# Big1 is a cell cycle regulator linking cell size to basal body number

**DOI:** 10.1101/2025.07.24.666660

**Authors:** Alexander J. Stemm-Wolf, Adam W. J. Soh, Lisa E. Mitchell, Huangqingbo Sun, Erik Collet, Gayatri L. Dholakia, Vikram Raju, Jay R. Hesselberth, J. Matthew Taliaferro, Robert F. Murphy, Lydia R. Heasley, Chad G. Pearson

## Abstract

Cell size control in dividing cells coordinates cell growth with cell division. In the ciliated protozoan, *Tetrahymena,* there is a tight link between cell size and the cytoskeletal assemblies at the cell cortex organized around basal bodies (BBs). BBs dictate the distribution of ciliary units governing cell motility and are organized into 18-22 ciliary rows. The number of BBs per cell remains remarkably consistent even when the number and lengths of ciliary rows vary. *big1-1* mutant cells are large and have elevated numbers of BBs, providing a system to investigate links between BB number and cell size control. We discovered *BIG1* encodes a protein with an RRM3 RNA-binding domain similar to the fission yeast meiotic entry gene, *mei2*. The *big1-1* mutation is a predicted null allele. By extending the duration of specific cell cycle stages conducive to new BB assembly, *big1-1* promotes cell size increases through BB amplification. In contrast, excess Big1 protein localizes to BBs and drives cells into premature cell division, resulting in small cells with fewer BBs. Thus, *Tetrahymena* Big1 localizes to BBs and controls cell cycle progression, indicating BBs and Big1 link cell growth to the cell division cycle.

## Background and Results

Cell size control in dividing cells operates through coordination between cell growth and division^1^. Cells must integrate information from their environment and from internal assessments of cell size and organelle assembly before committing to cell division. Extracellular conditions are transduced by signaling cascades and integrated with internal cell size signals to regulate entry into the cell cycle^2–6^. Internal control of a threshold size that is suitable for division can operate through either the dilution of negative regulators^7–9^ (proteins whose concentration decreases as cells grow) or the accumulation of positive regulators^10,11^ (proteins whose concentration increases during growth)^2,12^. Commitment to cell division is the fundamental execution point modulated by cell size controllers^1^. Cell division is coordinated by cyclin dependent kinases (CDKs) and their associated cyclins^13,14^. The regulation of Cyclin/CDKs is governed both by their abundance, via transcriptional control^15,16^, and by their activity, which is modulated by non-coding RNAs^17^, RNA-binding proteins^18,19^, inhibitory proteins, phospho-control of protein activity^20,21^, and proteasomal degradation^22,23^.

Spatial regulation of cyclin/CDK activity is an important yet incompletely understood mechanism of cell cycle control. Cell cycle control is modulated both at the cell cortex and the centrosome, which serves as a signaling site that recruits regulatory proteins^24^ including CDK1/cyclinB^25–28^, CDK2/cyclinE^29^, PLK1^30,31^, Aurora kinase^32^, CDC25^33^, and Wee1^34–36^. At the centrosome core are evolutionary ancient structures called centrioles that have a second role at the cell cortex where they act as basal bodies (BBs) to organize cilia. BBs may also exert an influence on cell cycle regulation^37,38^. The cell cortex itself regulates cell size. For example, *S. pombe* Cdr2 accumulates in cortical nodes at the middle of the cell to promote cell division based on cell surface area^11,39,40^. Together, these observations suggest that basal bodies may integrate regulatory signals at the cortex to coordinate cell cycle progression with cell size control.

In metazoans, terminally differentiated multiciliated cells produce hundreds of centrioles that act as BBs to nucleate cilia at the apical surface for fluid flow. There is a tight correlation between the number of BBs and apical surface area, but whether BB number or surface area control cell size has yet to be fully resolved^41–43^. The total number of BBs in the ciliate, *Tetrahymena,* is tightly controlled through the cell cycle^44–46^. Additionally, *Tetrahymena* CDK1 localizes to BBs^37^. This suggests an evolutionarily ancient centrosome/BB integrative function for information important to cell division and cell size control. Here we utilize the *Tetrahymena big1-1* mutant, which results in large cells with a concomitant increase in BBs, to understand the connection between the cortical cytoskeleton, the cell cycle, and cell size control.

### The *big1-1* mutant produces excess basal bodies, increasing cell size

The *big1-1* mutant was isolated in a genetic screen of *Tetrahymena* following nitrosoguanidine mutagenesis and results in large cells with extra ciliary rows (Figure1A)^47^. To determine whether the increase in cell size resulted from increased BB numbers, BBs (Poc1-mCh) were quantified using an automated imaging processing pipeline^44,48^. BBs, BB assembly, and cell geometry were measured in WT and *big1-1* cells. In addition to being longer and wider than WT cells (Supplemental Figure 1A), *big1-1* cells have more cortical rows with more BBs in each row (Supplemental Figure 1B).

To determine whether excess BBs promote increased cell size or whether an increased cell size induces BB assembly as a space filling measure, surface area (SA), volume (V), and BB assembly were measured in cycling *big1-1* cells. A greater than 2-fold excess in V and BB number was observed in *big1-1* (Figure 1B, Supplemental Figure 1B). To identify the causative change, BB number was plotted against V. The slope of the linear best fit decreased in *big1-1*, indicating BB number increases before volume in *big1-1* cells (Figure 1C). If BB number increases before V then the spacing between BBs (the inter-BB distance) should also be reduced in *big1-1* cells. Indeed, this was observed, particularly in the medial and posterior regions of the cell (Figure 1D). To identify the most significant differences between WT and *big1-1* cells, eleven measured properties of each cell were subjected to principal component analysis (see Methods; Figure 1E). Principal component 1 (PC1) contains 66.12% and PC2 16.47% of the variance in the data set and plotting the cells along these axes separates WT from *big1-1* (Figure 1E, Supplemental Figure 1C). PC1 is largely defined by primary geometric measurements of the cell (width, length, SA, V, inter-BB distance) all of which positively correlate with *big1-1* except for inter-BB distance which has a strong negative correlation. Thus, BB density is a defining characteristic of *big1-1* cells. In sum, these results support the conclusion that excess BBs are upstream of the increased cell size observed in *big1-1* and that increased BB assembly promotes enlarged cell size.

**Figure 1.**
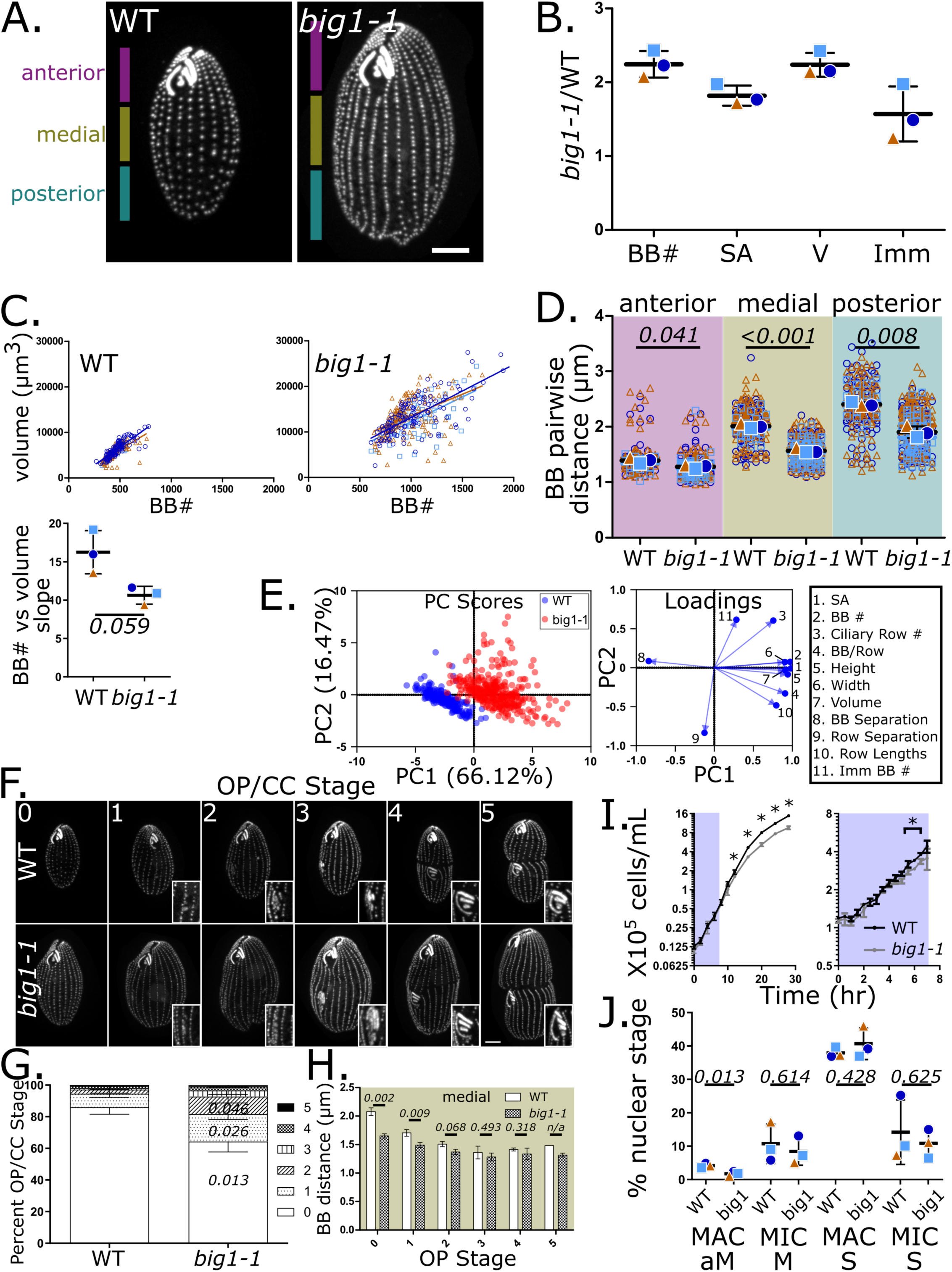
A dysregulated cell cycle leads to increases in BB number and cell size in *big1-1.* A) A wild type (WT) and *big1-1* cell visualized by Poc1-mCherry. Anterior, medial and posterior thirds of the cells are delineated. B) *big1-1*: WT ratios of basal body number (BB), surface area (SA), volume (V) and immature basal body number (Imm). C) BB number plotted against volume. The linear best-fit curves for each replicate are plotted on the graphs as well as in the graph below, showing volume cannot keep pace with BB number. D) The distance between BBs in the anterior, medial and posterior regions of cells. E) A principal component analysis (PCA) using the 11 inputs identified in the box. Loadings indicate the makeup of PC1 and PC2. F) WT and *big1-1* were staged based on their overall morphology and the development of the oral primordium (OP). G) Percentage of cells at each stage delineated in F). G) The inter-BB distance in the medial region of cells separated by cell cycle and OP stage. I) Growth curves of WT and *big1-1* cells grown at 30°C. The growth curve on the left shows 30 hours of growth with cells counted every two to four hours. The growth curve on the right shows seven hours of growth with cells counted every ½ hour. “*” indicates a p value <0.05. J) Nuclei in WT and *big1-1* with mitotic morphologies or in S phase by incorporation of EdU after 20 minutes of labeling. Scale bars = 10 µm. Error bars show standard deviations of three biological replicates.

To directly test whether there is more BB assembly in *big1-1* mutants, we examined the number of new, immature BBs (imm BB). The absolute number of imm BBs is elevated in *big1-1* (Figure 1B); however, the proportion to total BBs is reduced (Supplemental Figure 1D). This suggests that BB assembly in *big1-1* occurs at a reduced rate, or that BBs mature more quickly in *big1-1*. To address this, BB maturity was quantified both by maturation of Poc1-mCh levels and by the ability to nucleate a cilium. These parameters were measured as a function of distance from the mother BB, a proxy for BB age^48–50^. In *big1-1*, both Poc1-mCh levels were elevated and cilia were present in daughter BBs near to mothers suggesting that BB maturation is accelerated (Supplemental Figure 1E). This interpretation is complicated by the decreased BB spacing in *big1-1*. However, increased maturation was observed even in close proximity (< 1.4 µm; Figure 1D) to mother BBs, leading us to conclude that BBs mature more rapidly in *big1-1* cells. Finally, to address whether more BB assembly occurs in *big1-1*, cells were arrested by media starvation to generate a uniform population of G1 cells with mature BBs. Cells were then released by media addition and BB assembly was quantified. *big1-1* cells assembled BBs more rapidly than controls, suggesting BB assembly is accelerated (Supplemental Figure 1F). In summary, *big1-1* mutants exhibit elevated BB maturation and assembly that causes enlarged cells by increasing cell length, width, SA, and V.

### *big1-1* mutants delay the cell cycle at phases associated with basal body assembly

One way in which BB assembly might be increased in *big1-1* mutants is through a delay during cell cycle phases that promote BB assembly. During *Tetrahymena* cell growth, a new oral apparatus, called the oral primordium (OP), is formed in the medial region of the cell. This serves as a morphological marker for cell cycle staging (Figure 1F). Most BB assembly during the cell cycle occurs at early stages of OP development (stages 0-2)^45,51^. In contrast to WT, the fraction of *big1-1* cells with a developing OP, particularly in stages 1 and 2, is significantly increased (Figure 1G). Furthermore, the inter-BB distance is significantly reduced in *big1-1* between stages 0 and 3 of the cell division cycle (Figure 1H), indicating these stages of the cell cycle are when BB assembly outpaces cell growth in *big1-1*. This suggests *big1-1* cells delay in a cell cycle phase that promotes continuous BB assembly. WT cells also produce more BBs at these phases of the cell cycle but progress more rapidly through them.

The increase in stage 1-2 *big1-1* cells suggests that either the total cell cycle time is elongated, or that other phases of the cell cycle shorten to compensate. Although the *big1-1* growth rate is reported to be 25 to 50% slower than WT controls^47^, we observed only a mild growth delay during logarithmic growth (Figure 1I left and right panels). We conclude that in logarithmic growth, *big1-1* compensates for slowed early phases of the cell cycle by more rapid progression through the cell cycle at other phases^52–55^. An analysis of DNA synthesis and nuclear division shows that macronuclear (MAC) and micronuclear (MIC) S phases are unaffected in *big1-1* mutants. However, there is a reduction in the frequency of cells undergoing MAC amitosis which occurs late in the cell cycle coincident with cytokinesis (Figure 1J)^56,57^. This supports the conclusion that *big1-1* compensates for prolonged stages 1-2 of the cell cycle with rapid passage through either early (stage 0) or late (stage 5) stages. Thus, the increased BB assembly and large cell phenotype of *big1-1* is likely driven by a cell cycle delay during phases conducive to BB assembly.

### The *BIG1* gene encodes a Mei2-like RNA-binding domain protein

Next-generation short read sequencing was used to identify the mutation responsible for the *big1-1* phenotype. Insertions were discovered in genes annotated as *RRM51* and *DPK1* in *big1-1* DNA (Figure 2A). Despite being a homozygous mutant, several distinct variant alleles of these disruptions were identified in the sequencing data, indicating the presence of an underlying genetic instability at these loci resulting in the development of a mosaic mutant state. For all alleles of the *RRM51* gene, the open reading frame was invariably disrupted in *big1-1*. This was not the case for *DPK1*; several alleles of the insertion affecting this gene preserved the open reading frame. *RRM51* resides on MIC chromosome 3 while *DPK1* resides on MIC chromosome 4. Crosses to MIC nullisomic lines determined that the gene responsible for the *big1-1* phenotype is on chromosome 3 (Supplemental Figure 2A). PCR across the predicted *BIG1/RRM51* breakpoint and insertion site produced the expected 297 base pair product from WT genomic DNA. However, many high molecular weight bands (1kb-7kb) and no 297 base pair band were observed in *big1-1*, indicating multiple distinct DNA species are inserted into this site. Sequencing of purified mutant species demonstrated that all alleles shared a common insertion site in the *RRM51* gene and that the variable insertion sequences were derived from a shared intergenic region on MIC chromosome 5 (Supplemental Figure 2B). This indicates that the *big1-1* phenotype arises from disruptions of *BIG1*/*RRM51* (hereafter *BIG1*).

**Figure 2.**
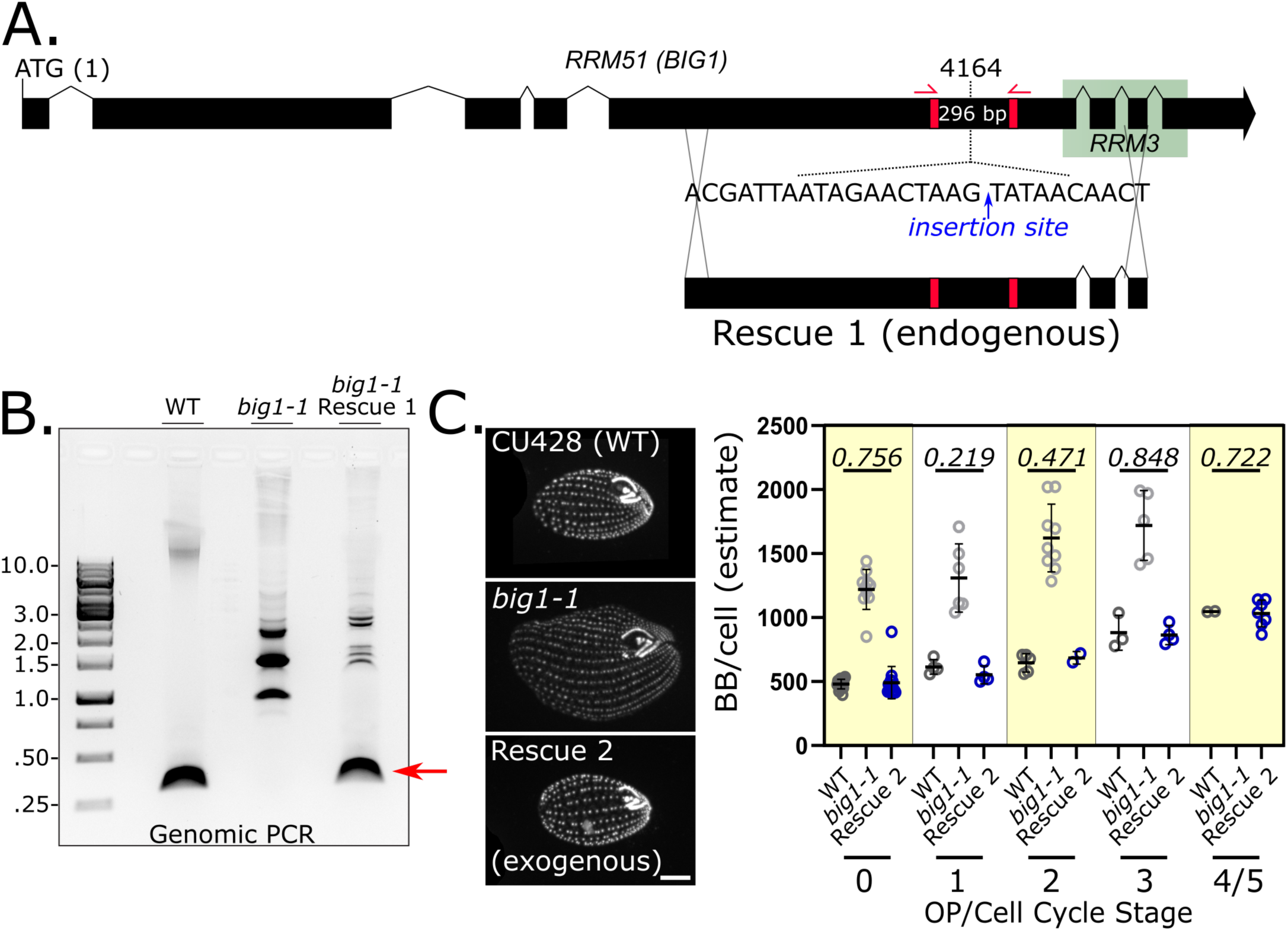
Identification of the *BIG1* gene. A) A schematic of *RRM51* showing the site of insertion in *big1-1*. The rescuing fragment (Rescue 1, a repair of the endogenous locus) is shown below. Red bars and arrowhead indicate the sites of primers designed to flank the insertion site. B) An agarose gel showing PCR products from DNA isolated from WT, *big1-1* and *big1-1*+Rescue 1. The red arrow denotes the expected size of the WT fragment. C) Images of a WT strain (lab strain CU428), *big1-1* and Rescue 2 (the entire RRM51 gene with *cis* regulatory elements cloned into an exogenous site) stained with an antibody to the BB marker, centrin. BB counts show that the BB numbers in the rescue strain are statistically similar to WT. Scale bar = 10 µm. Error bars show standard deviations.

Two approaches were taken to determine whether introducing WT *BIG1* into *big1-1* cells rescues the large cell phenotype. First, a WT DNA fragment spanning the insertion site was transformed into *big1-1* cells (Figure 2A). Cells with WT morphology were recovered and termed Rescue 1 cells. Second, an exogenous construct containing *BIG1* and *cis* regulatory elements fused at the N-terminus to an mCherry (mCh)-6XHIS tag was transformed into *big1-1* cells (Supplemental Figure 2C). Transformants (Rescue 2) displayed WT morphology and restored BB number to WT levels (Figure 2C). PCR from rescued cells restored the expected 297 base pair fragment, although some mutant species remained, indicating mutant alleles are recessive (Figure 2B). Because *big1-1* cells have insertions in *DPK1*, the endogenous *BIG1* rescue (Rescue 1) was used as the control strain for *big1-1* in the automated image analyses (Figure 1, Supplemental Figure S1).

*BIG1* encodes a 1473 amino acid protein with an RNA recognition motif most similar to the RRM3 domain of the fission yeast *mei2*, which is essential for entry into meiosis^58,59^. Mutations in related plant genes, *OML4* and *AML1* cause large rice grains due to larger spikelet hull cells in the former^60^ and early *Arabidopsis* bolting or stem elongation in the latter^61^. Thus, this conserved protein plays known roles in major cell cycle transitions and cell size control. Attempts to localize Big1 using Rescue 2 by mCh fluorescence, antibodies to mCh, or antibodies to 6XHIS were all unsuccessful, suggesting that protein expression is below our detection threshold. However, *BIG1* RNA expression was observed (Supplemental Figure 2D). In summary, we identified the gene responsible for the *big1-1* phenotype to be an RNA-binding domain containing protein analogous to other cell cycle and cell size regulators.

### *BIG1* expression accelerates the cell cycle and decreases cell size

To test whether *BIG1* expression has the inverse effect as the *big1-1* phenotype, an inducible mCh-6HIS tagged *BIG1* expression construct was introduced into WT cells^62,63^. After 5 hours of *BIG1* overexpression, small cells were observed in the cell population (Figure 3A). Next, a time course of *BIG1* overexpression was performed. Ratio analyses between *BIG1* induced and uninduced cells revealed that the number of immature BBs was the first change detected in these cells. Significantly fewer immature BBs were observed 1 hour after *BIG1* induction (Figure 3B, Supplemental Figure 3A). Thus, elevated Big1 quickly reduces BB assembly. Cell growth during the time course revealed a short, transient acceleration in doubling time between 2 and 3 hours, indicating *BIG1* overexpressing cells divide before their uninduced counterparts, and that *BIG1* overexpression accelerates the cell cycle (Figure 3C, Supplemental Figure 3B).

**Figure 3.**
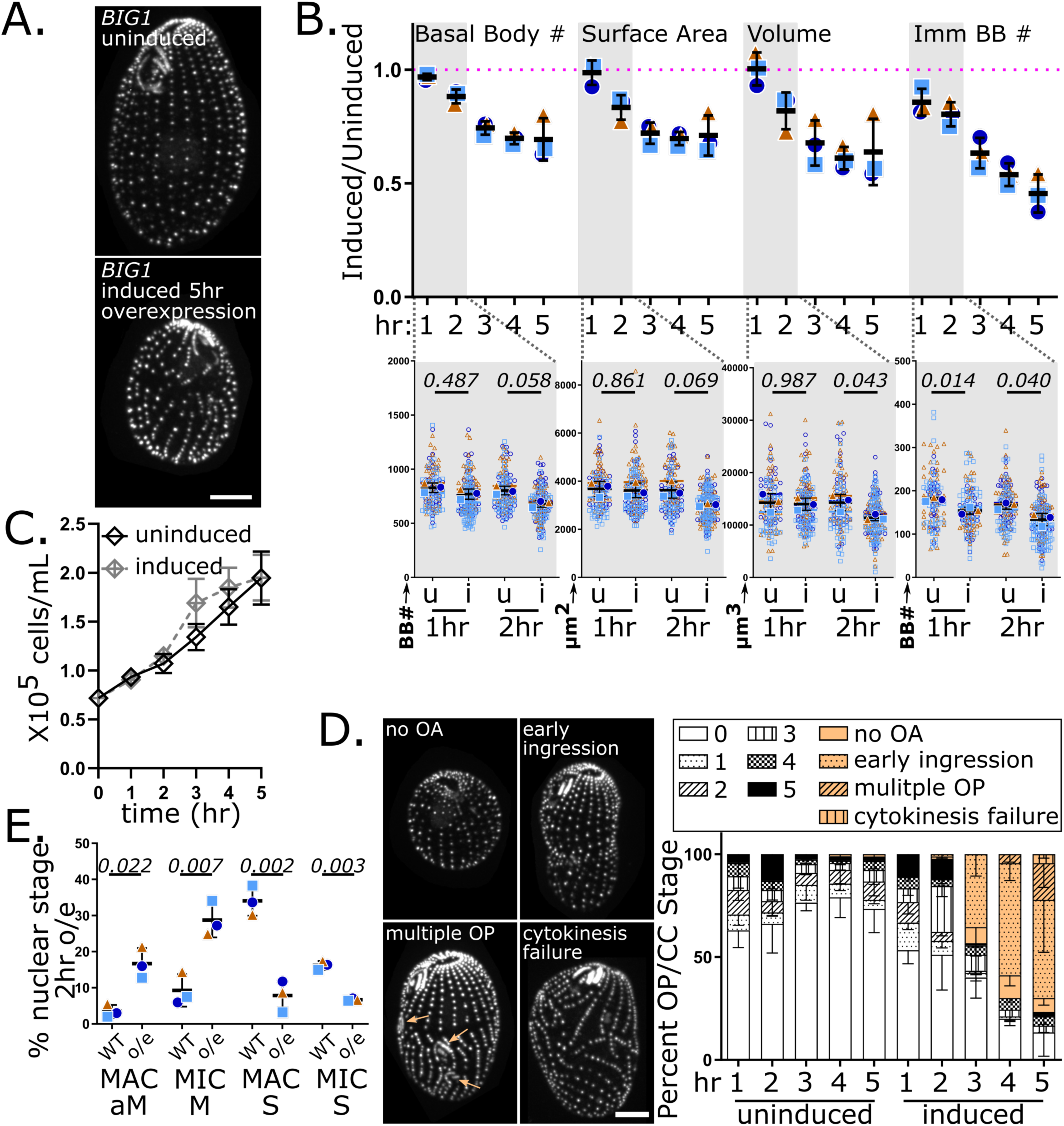
Overexpression of *BIG1* leads to an accelerated cell cycle and smaller cells with fewer BBs. A) Images of a strain harboring a cadmium driven promoter integrated in front of the *BIG1* gene in the absence (top) or presence (bottom) of cadmium. BBs are visualized by Poc1-GFP. B) Induced: Uninduced ratios through a five-hour time course of *BIG1* induction. The magenta line indicates a ratio of 1. The gray shading indicates the first two hours of the time course, and the full data is shown below. u = uninduced, i = induced. C) Growth curves from the cultures used in the time course. D) Examples of aberrant morphologies observed with *BIG1* overexpression. The percentage of each OP, cell cycle stage and aberrant morphology is plotted to the right. Orange arrows indicate the formation of excess OPs. E). Nuclei in *BIG1* uninduced and induced cells with mitotic morphologies or in S phase by incorporation of EdU after 20 minutes of labeling. Scale bars = 10 µm. Error bars show standard deviations of three biological replicates.

To determine the timing of progress through the cell cycle, cells were staged based on OP development and overall cell morphology. As the *BIG1* overexpression time course progressed, fewer stage 0 cells were observed giving way to more late division stage cells (stages 3 to 5) (Figure 3D). The nuclear cell cycle was also severely affected after 2 hours of *BIG1* overexpression with significant increases in the number of cells in MAC amitosis and MIC mitosis, and reductions in both MAC and MIC DNA synthesis (Figure 3E, Supplemental Figure 4A). After 3 hours of *BIG1* overexpression, abnormalities were detected, most notably early cytokinetic furrow ingression without sufficient OP development, and additionally cells with multiple OPs, missing oral apparatuses (OA) (presumably due to cytokinesis without a developed OP), and cells that had aborted cytokinesis (Figure 3D). In summary, overexpression of *BIG1* inhibits BB assembly and accelerates the cell cycle, resulting in small cells with fewer BBs and cell morphology defects.

### Elevated *Big1* overrides starvation induced cell cycle arrest

External nutrients activate signaling networks promoting growth and entry into the cell cycle^64^. In the absence of nutrients, these networks are inactivated by GCN2 signaling which reduces translation and arrests cells prior to cell cycle commitment^65–68^. Starvation of *Tetrahymena* arrests cells in G1 at OP stage 0^69^. To investigate how Big1 interacts with this pathway, *BIG1* was overexpressed in starvation arrested cells. Whereas uninduced cultures remained at the same cell density, *BIG1* overexpressing cell density increased slightly after 3 hours (Figure 4A). After 4 hours of *BIG1* overexpression, 38.5% of the cells had either assembled an OP, begun cytokinetic furrow ingression, or had gone through cell division with an incomplete OP. In contrast, uninduced cells were invariably in stage 0, as expected for starvation arrested cells (Figure 4B, E). Thus, *BIG1* overexpression subverts normal starvation-induced cell cycle arrest, causing cells to enter into cell division, producing morphological defects.

**Figure 4.**
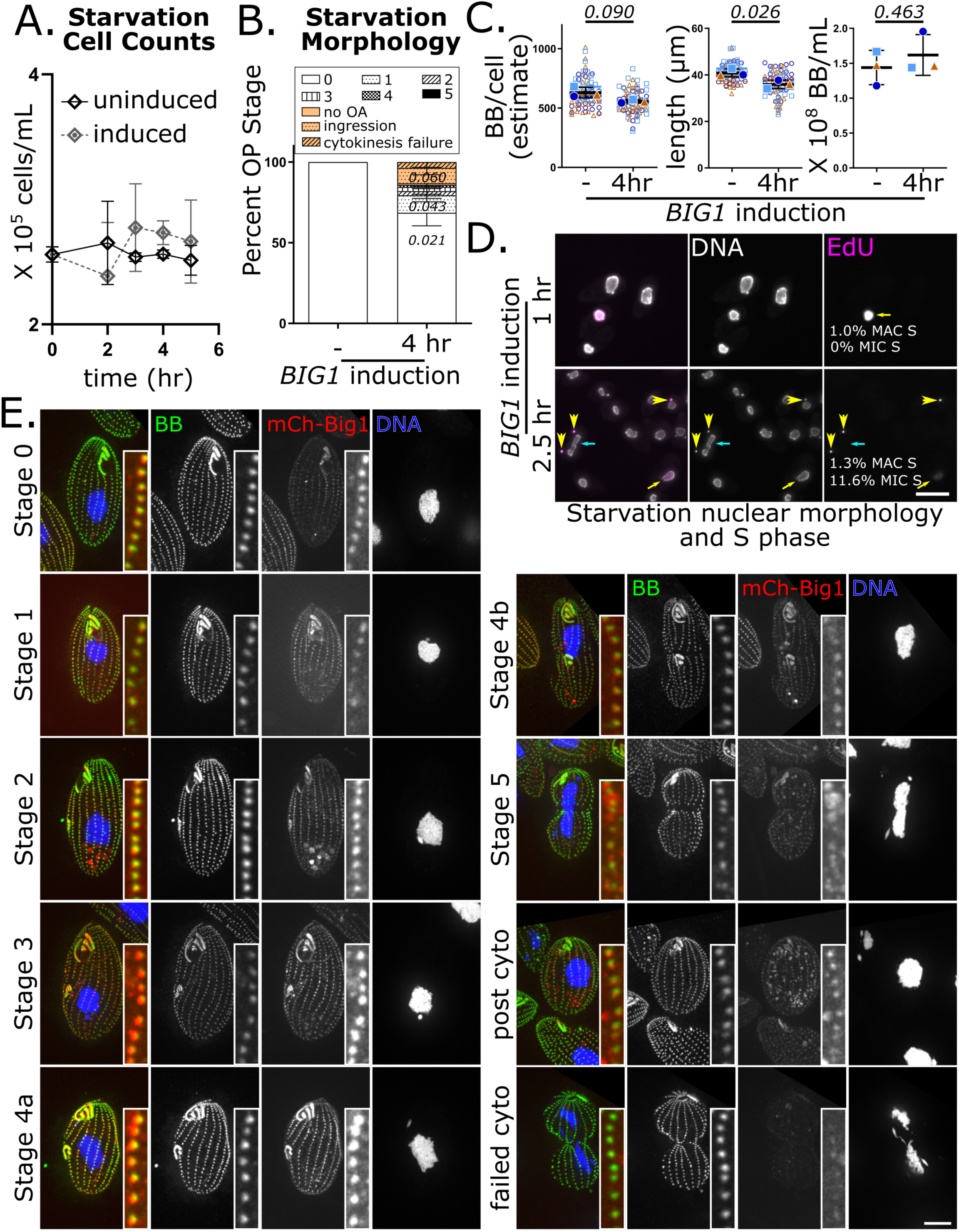
Elevated Big1 causes starvation arrested cells to enter the cell cycle and colocalizes with BBs. A) Cell counts of starvation arrested cells after induction of *BIG1*. B) The percentage of cells showing OP assembly and cell cycle progression in *BIG1* uninduced and 4 hour induced conditions. Aberrant morphologies were observed and included in the analysis. C) Quantification of BB number, cell length and the total number of basal bodies in *BIG1* uninduced and 4 hour induced conditions. D) Starvation arrested cells were induced for *BIG1* expression and S phase was monitored by EdU incorporation 1 and 2.5 hours after induction. Yellow arrows indicate MAC S phase, yellow arrowheads indicate MIC S phase and the cyan arrow indicates a MAC going through amitosis. Scale bar = 50 µm. E) Localization of overexpressed mCherry-Big1in starvation arrested cells. Insets show colocalization with the centrin BB marker. Scale bar = 10 µm and insets are 2 µm X 10 µm. Error bars show standard deviations of three biological replicates.

To address whether BBs assemble in starvation arrested Big1 overexpressing cells, a count of the number of cortical BBs per cell was generated and multiplied by the cell density to estimate the total number of BBs/mL. This calculation factors out any reduction in BBs per cell due to cell division. No significant change in cortical BB number was detected (Figure 4C). This suggests that BB assembly is sensitive to nutrient availability and could act as an independent measure of cell size. Consistent with some of the cells having gone through cell division, the average length of *BIG1* overexpressing cells was significantly reduced (Figure 4C). Thus, elevated Big1 induces arrested cells to enter the cell cycle without assembling new BBs.

To address whether *BIG1* overexpression-induced entry into cell division promotes nuclear events of the cell cycle, DNA synthesis and nuclear division were quantified at 1 and 2.5 hours after *BIG1* overexpression. In normal cycling *Tetrahymena* cells, MAC S occurs prior to MIC S^70^. After 1 hour of *BIG1* overexpression, 1.0% of the cells were in MAC S and no cells were in MIC S; after 2.5 hours, 1.3% of cells were in MAC S and 12.0% in MIC S (Figure 4D and Supplemental Figure 4B). Uninduced cells were never observed in either S phase. Therefore, *BIG1* overexpression induces cells to replicate both MAC and MIC DNA. Observation of MAC S was rare. Because MAC S occurs earlier in the cell cycle than MIC S^70^, it suggests that cells are more often at late stages of the cell cycle during *BIG1* overexpression. After 2.5 hours of induction, 3.0% of the cells were observed in MAC amitosis and 5.0% in MIC mitosis (Supplemental Figure 4B). Cells in MAC and MIC amitosis and mitosis, respectively, continued to be observed 4 and 5 hours into overexpression (Figure 4E). In conclusion, overexpression of *BIG1* is sufficient to override starvation induced cell cycle arrest and commit cells to the division cycle, including DNA synthesis, but in the absence of BB assembly. This creates a reductional division of BBs and smaller cells.

### Elevated Big1 localizes to basal bodies

Cell cycle machinery is commonly associated with centrosomes and BBs^24^. To investigate where Big1 influences the cell division cycle, we explored its localization. Big1 colocalizes with the BB marker, centrin, at BBs^71^ (Figure 4E). At early stages of the cell cycle, two gradients of BB localization were observed: first is a gradient along the anterior-posterior axis with the majority of Big1 localizing to the medial region of the cell; second is a lateral gradient with a strong signal along the cortical rows immediately beneath the oral apparatus (row 1 and its immediate neighbor to the cell’s left). Intriguingly, these gradients could define the placement of a new OP and overlap with the gradients of Hippo signaling molecules that assure proper cytokinetic furrow placement^38,72,73^. As cells progress into later stages of the cell cycle, the Big1 signal intensifies and expands to more BBs such that gradients are less apparent. When cytokinetic furrow ingression begins, Big1’s BB signal declines. With the caveat that we could only localize overexpressed *BIG1*, we conclude that Big1 localizes to BBs and that BBs are key structures in regulating cell size through control of the cell cycle.

## Discussion

Cell size control across species is thought to optimize cell function and fitness^1^. In *Tetrahymena*, this is reflected in their swimming speeds which are adversely affected when cells are big (*big1-1*) or small (elevated Big1) (Supplemental Figure 4C). We discovered that Big1 is a major negative regulator of cell size through its role in promoting cell cycle entry. Big1 localizes to BBs, suggesting BB number may be how *Tetrahymena* cells monitor cell size and surpassing a threshold BB number commits *Tetrahymena* to cell division. This model accounts for the strict constancy of BB number, even with variation in the number of BB cortical rows or their lengths^46^. The mechanisms underlying how Big1 promotes cell cycle progression remain to be elucidated.

Several lines of thought can be applied to potential mechanisms of Big1 function. The related RNA-binding protein, *S. pombe* Mei2, is required for entry into meiosis, operating through the sequestration of a meiotic inhibitor^58,74–76^. The potential sequestration of a cell cycle inhibitor at BBs through Big1 would tie BB number to commitment to cell division. This model invokes a constant level of a cell cycle inhibitor that is progressively sequestered at BBs, such that, as BB number increases, a threshold of inhibitor depletion from the cytoplasm commits cells to division. An alternative, yet equally plausible model, is Big1 facilitated activation of cell cycle regulators at BBs that, once activated, commit cells to division. This is consistent with the presence of cell cycle activators at BBs, including CDK1^37^ and the polo-like family of kinases (PLKs)^77^. Furthermore, mitotic regulators localize to and are activated at centrosomes in other organisms^26,30–36,78–80^.

Big1 itself is regulated through the cell cycle. Initially Big1 localization is defined by a gradient along the anterior-posterior axis and along a lateral axis (Figure 4E). This localization occurs prior to the development of an OP. MAC S phase also occurs prior to OP development^51,70^, yet its occurrence suggests the satisfaction of a restriction point committing cells to division. Given that Big1 overexpression can induce starvation arrested cells to enter S phase (Figure 4D), and Big1 is largely restricted to the two rows immediately posterior to the OA at this time, one might speculate that S phase entry is sensitive to the number of BBs in these rows. Previous work shows a 1.5 – 2-fold increase in BB number during stage 0 in these two cortical rows^44^. Once cells commit to MAC S, Big1 localization spreads to the other BBs in the cell, perhaps regulated by S phase CDKs. Big1 contains several CDK sites, including a canonical S/T-P-x-R/K (SPEK) site N-terminal to the RRM3 RNA-binding domain, and S phase cyclins are known to localize to centrosomes^29,38,81,82^. This higher level of BB localization may constitute a second trigger that facilitates entry into mitosis. Interestingly, the *Chlamydomonas* RNA-binding protein encoded by *TNY1* acts at a G1 size control point which commits cells to division once a size threshold has been achieved and at the S/M control point which determines the number of divisions cells go through based on size^19^, suggesting a conserved role for RNA-binding proteins in cell size control.

If Big1 is a critical regulator of cell cycle timing, how do *big1-1* cells manage to divide with a growth rate similar to that of WT cells? One possibility is the presence of other genes with the RRM3 domain (there are three in *Tetrahymena*) which could potentially compensate for the loss of *BIG1*. If the RRM3 domain proteins bind overlapping sets of RNAs, *BIG1* overexpression would have a global effect in RNA binding/sequestration, which may explain the dramatic effects on growth rate and cell cycle morphology observed with *BIG1* overexpression (Figure 3C, D and Supplemental Figure 3B).

In summary, we find the RRM3 domain-containing *BIG1* gene is a cell cycle timing regulator that maintains cell size. Its localization to BBs suggests that cell size control is tightly linked to BBs and that BB number expansion may trigger commitment to cell division.

## Methods

### Cell lines and culture conditions

Strains used in this study (*big1-1* [IA313], IA267, CU428 and micronuclear nullisomic lines) were from the Tetrahymena Stock Center (Washington University, St. Louis, MO, formerly Cornell University, Ithaca, NY). An isogenic *BIG1* strain (WT) was made by transforming the *big1-1* cell line with a rescuing fragment that spans the *big1-1* insertion site. p4T2-1-Poc1-mCh-NEO2 conferring paromomycin resistance was transformed into *big1-1* and WT. *BIG1* overexpressing lines were made with p4T2-1-MTTpr-mCh-*BIG1*, which confers paromomycin resistance. This construct was transformed into IA267 and into IA267 containing Poc1-GFP-BSR (blasticidin resistance). Cells were grown in SPP and starved in 10 mM Tris, pH 7.4^83^. Cell counts were performed with a Z1 Coulter Counter (Beckman Coulter, Brea, CA). Transformations were performed using a PDS-1000 biolistic particle delivery system with 900 psi rupture disks and 0.6 µm Tungsten particles (Bio-Rad, Hercules, CA). Inducible genes were under the control of the *MTT1* promoter^63^ and induced using 0.5 µg/mL CdCl_2_ in SPP and 0.1 µg/mL CdCl_2_ in 10 mM Tris, pH 7.4.

### Constructs

p4T2-1-Poc1-mCh was generated by cloning 1 kilobase flanking regions into pmCherryLAP-NEO2^62^ to place the mCh tag at the 3’ end of *POC1*. For p4T2-1-Poc1-GFP-BSR, *POC1* flanking sequences as described above were cloned into p4T2-1-GFP-NEO2 followed by the replacement of the NEO2 cassette conferring paromomycin resistance with the BSR cassette conferring resistance to blasticidin^84^. The *BIG1* overexpression construct was made by cloning one kilobase flanking regions present upstream and in the 5’ gene region into pNEO2-MTT1pr-mCherryLAP^62^. The exogenous Rescue 2 construct was cloned into pBS-RPL29 vector conferring cycloheximide resistance (the kind gift of D. Chalker, Washington University, St. Louis, MO). First, an 1100 base pair region containing the *BIG1* promoter was cloned in, followed by a region containing mCh-6HIS and some of the 5’ region of the *BIG1* gene from the p4T2-1-MTTpr-mCh-*BIG1*. The rest of the gene, including 667 base pairs of downstream sequence was subsequently cloned in. All oligonucleotides used in this study are listed in Supplemental Table 2.

### Fluorescence and immunoflourescence microscopy

For automated image analysis, WT and *big1-1* with Poc1-mCh, and IA267/p4T2-1-MTTpr-mCh-*BIG1*/Poc1-GFP were stained with 1:500 dilutions of an antibody to mCh (Living Colors mouse monoclonal antibody, Takara Bio USA, San Jose, CA) or an antibody to EGFP^85^ (rabbit polyclonal, the kind gift of Michael Rout, the Rockefeller University, New York, NY), respectively. 750,000 – 1,000,000 mid-log phase cells were washed in 10 mM Tris and fixed for 5 minutes in PHEM with 3.2% Paraformaldehyde and 2.4% Triton X-100^86^. Cells were pelleted at 600g for 1 minute and permeabilized for 10 minutes in ice-cold PHEM with 0.5% Triton X-100. Cells were then washed 3X PHEM with 0.5% BSA. Primary and secondary antibody incubations were carried out at room temperature for 1.5 hours and 1 hour, respectively, diluting antibody into PHEM with 0.5% BSA. Secondary antibodies were anti-Mouse Alexa 594, and anti-Rabbit Alexa 488 (Thermo Fisher, Waltham, MA), diluted 1:1000. Cells were washed 3X with PHEM with 5% BSA after each antibody incubation. 10 µl of the final pellet was mixed with 90 µl antifade (1X PBS, 90% Glycerol, 0.5% n-Propyl Gallate) and put into a homemade chamber using cover glass affixed to a slide using double-stick tape and then sealed with VALAP (1:1:1 Vaseline, lanolin and paraffin). Slides were incubated overnight at 4°C with the cover glass down to allow gravity to pull cells closer to the cover glass prior to imaging. For Centrin and DNA staining with mCh-*BIG1* autofluorescence imaging, 750,000 – 1,000,000 cells were spun down, washed in PHEM, and fixed for 20 minutes in PHEM with 3.2% paraformaldehyde and 0.24% Triton X-100 at room temperature. 10 mL of PBS with 0.1% BSA was added prior to 3 washes with PBS with 0.1% BSA. Cells were stored in PBS with 0.1% BSA at 4°C until antibody processing. 15 µl of cells were dried onto a cover glass, then blocked with PBS with 1% BSA for 30 minutes. 1:2000 dilution of primary antibody (anti CEN1 2111^71^) and 1:1000 dilution of secondary antibody (anti Rabbit Alexa 488 mixed with Hoechst 33342 (Thermo Fisher)) were for 1.5 and 1 hour respectively at room temperature, followed by three washes in PBS with 0.1% BSA. Coverslips were mounted on antifade, sealed with clear nail polish and imaged. For EdU staining, cells were pulsed for 20 minutes with 25 µM EdU. Cells were then washed twice with PBS and fixed in PBS with 4% paraformaldehyde for 15 minutes. Cells were washed twice with PBS prior to overnight storage. 15 µl of cells were dried on cover glass, and then permeabilized for five minutes in PBS with 0.5% Triton X-100 followed by two PBS washes. A mixture containing 0.2 µM AZDye Picolyl Azide (Cy5 picolyl azide, Vector Laboratories, Newark, CA), 0.8 mM CuSO4 and 1 mg/mL Sodium L-ascorbate (made fresh) in PBS was added and incubated, protected from light, for 30 minutes^70,87^. After a PBS wash, cells were counterstained with Hoechst 33342 for five minutes, washed with PBS, mounted on antifade and sealed with clear nail polish.

Samples were imaged on a Nikon Eclipse Ti inverted microscope stand equipped with a 60X Plan Apochromat oil objective (NA 1.40) and a 100X Plan Apochromat oil objective (NA 1.45) (Nikon Instruments, Inc., Melville, NY), an Andor iXon X3 camera (Oxford Instruments, Abingdon, UK), and a CSU-X1 spinning disk confocal (Yokogawa Electric Corporation, Sugar Land, TX). For automated image analysis, the 60X objective was used along with a 0.2 µm step size. Imaging for EdU staining was done on a Nikon Eclipse Ti inverted microscope stand equipped with a 40X Plan Fluor dry objective (NA 0.75) equipped with a Teledyne Photometrics Kinetix 22 camera (Teledyne Vision Solutions, Billerica, MA). Slidebook 2023 digital microscopy software (Intelligent Imaging Innovations, Inc, Denver, CO) was used for image acquisition.

### Image analysis

For the automated image analysis, cells from images were manually segmented such that each image to be analyzed had only one cell. The TetAlize analysis software described in Sun et al^44^ was used to produce the results in Figures 1 and 3 and Supplemental Figures 1 and 3. All BBs were detected and those having <0.5 the fluorescence intensity of the nearest neighbor were defined as immature BBs. All cells were automatically aligned along the anterior-posterior axis to distinguish anterior, medial and posterior BBs. Manual estimates of BB number in Figures 2 and 4 were made by manually multiplying the cell length, the number of ciliary rows and the average of BBs per µm (generated by counting BBs along 10 µm of the medial region in six rows for each cell). Image processing and manual analyses were done using FIJI^88^.

### Genomic analysis of the *big1-1* mutation

*big1-1* (IA313) and IA267 were sequenced on an Illumina NovaSEQ6000 by the University of Colorado Anschutz Medical Campus Cancer Center Genomics Shared Resource Core Facility (RRID:SCR_021984). All genomic analyses were performed using CLC Genomics Workbench (Qiagen, Redwood City, CA). Macronuclear (MAC) and micronuclear (MIC) references for *Tetrahymena thermophila* were downloaded from the *Tetrahymena* Wiki at tet.ciliate.org (Bradley University, Peoria, IL) (MAC: 1-upd-Genome-assembly.fasta; MIC: 2016_mic.genome). Paired WGS read datasets were trimmed, mapped to the MAC reference genome and subjected to variant detection analysis to identify homozygous de novo single nucleotide polymorphisms (SNPs) and structural variations. Variant calls were then filtered to identify variants present in *big1-1* but not the wild type IA267. Results were manually screened and validated. Homozygous disruptions of *RRM51* (*TTHERM_*00444880) and *DPK1* (*TTHERM_00203010*) were identified. To identify the nature of the insertional mutation, primers flanking the insertion breakpoints (5’-CTCAAGCTCCTCCTCCTCAC-3’ and 5’-GCTGTTACCTTATTAGCTTTGG-3’) in the *RRM51* gene were used to amplify the predicted insertion by PCR, using Q5 DNA polymerase (New England Biolabs, Ipswich, MA). An extension time of 10 minutes was used to enable capture of long variants of the predicted insertion. These higher molecular weight fragments were gel purified, pooled, and sequenced with Oxford Nanopore single molecule long read sequencing technologies (Plasmidsaurus, South San Francisco, CA). These long read data were then mapped to both the MAC and MIC reference genomes to identify the origins of the insertion sequences. All insertions mapped to a repetitive region on MIC chromosome 5 spanning portions of the internally eliminated sequence element annotated at position 25,013,413 – 25,011,223. The ends of this element contain long stretches of homology oriented on opposite strands. The insertions defined using long read sequencing contained a portion of this repetitive sequence. This precluded our ability to determine the precise origins of the disrupting insertion sequence.

### RT-PCR

RNA was isolated from cultures in mid-log phase. 1 µg of RNA was used as a template for first strand synthesis using the SuperScript IV First-Strand Synthesis System (Invitrogen/Thermo Fisher) and the reverse primer flanking the insertion site, described above as per the manufacturer’s instructions. Using this cDNA as a template and the insertion site flanking primers, PCR was carried out using the OneTaq DNA polymerase and buffer (New England Biolabs) as per the manufacturer’s instructions.

### Swimming assays

WT and *big1-1* cells at 30°C were grown to mid to late-log phase and diluted to 200,000 cells/mL, allowed one hour of recovery prior to imaging at 30°C. *BIG1* inducible overexpressing lines were treated similarly, induced with 0.5 µg/mL CdCl_2_ at 100,000 cells/mL and grown for 24 hours, then diluted to 200,000 cells/mL and similarly allowed one hour of recovery. Imaging chambers were made by affixing an 18 × 18 mm cover glass onto a 30 mm round 1.5 cover glass with double-stick tape and pre-warmed to 30°C. Imaging was conducted while incubating cells at 30°C using a Warner Instruments TC-324C automatic temperature controller (Harvard Apparatus, Holliston, MA). 200 images were captured at 50 msec intervals. Cells paths were traced using the segmented line tool in FIJI throughout their time in the field of view and the rate was calculated based on distance travelled divided by time elapsed.

### Statistical methods

Graphs show individual data points, the mean value and the standard deviation from each replicate. Statistics were calculated on the mean values from three replicates using Graphpad Prism 10 (Graphpad Software, Boston, MA), or Microsoft Excel (Microsoft, Richmond, WA). The comparison of WT to Rescue 2 used the Student’s t-tests performed on all data points. Student’s t-tests did not assume equal variation between different strains and conditions (Welch’s correction or heteroscedastic t-test). Equal variation was assumed between replicates for growth curves. Principal Component Analysis was done in Graphpad Prism 10 on 11 properties from the TetAlyze analysis, listed in Figure 1E and standardized to have a mean of 0 and standard deviation of 1. PCs were selected based on parallel analysis. A complete list of statistical tests and N values is in Supplemental Table 1.

## Resource availability

Reagents generated in this study will be made available on request. TetAlyze software is available from https://github.com/murphygroup/TetAlyze.

## Author contributions

Conceptualization, AJSW, AWJS, LEM, RFM, LRH, CGP; Investigation, AJSW, AWJS, LEM, EC, GLD, VR, JRH, JMT, LRH, CGP; Formal Analysis, AJSW, AWJS, LEM, HS, LRH, RFM; Writing – Original Draft, AJSW; Writing – Review & Editing, AJSW, CGP; Funding Acquisition, CGP.

## Acknowledgements

The authors gratefully acknowledge Joseph Frankel at the University of Iowa, Iowa City, IA for the isolation, early characterization and discussions regarding the *big1-1* mutant strain. Tenzin Dolma Olsen cheerfully and helpfully assisted with *BIG1* overexpression experiments. Members of the Pearson lab contributed insightful discussions.

CGP: R35GM140813

JRH: R35GM119550

LRH: R00GM134193

Cancer Center Support Grant: P30CA046934

## Declaration of interests

The authors declare no competing interests.

## Abbreviations

WT: wild type
BB: basal body
SA: surface area
V: volume
Imm BB: immature basal body
OA: oral apparatus
OP: oral primordium
PCA: principal component analysis
mCh: monomeric cherry

## Supplemental Figure Legends

**Supplemental Figure 1.**
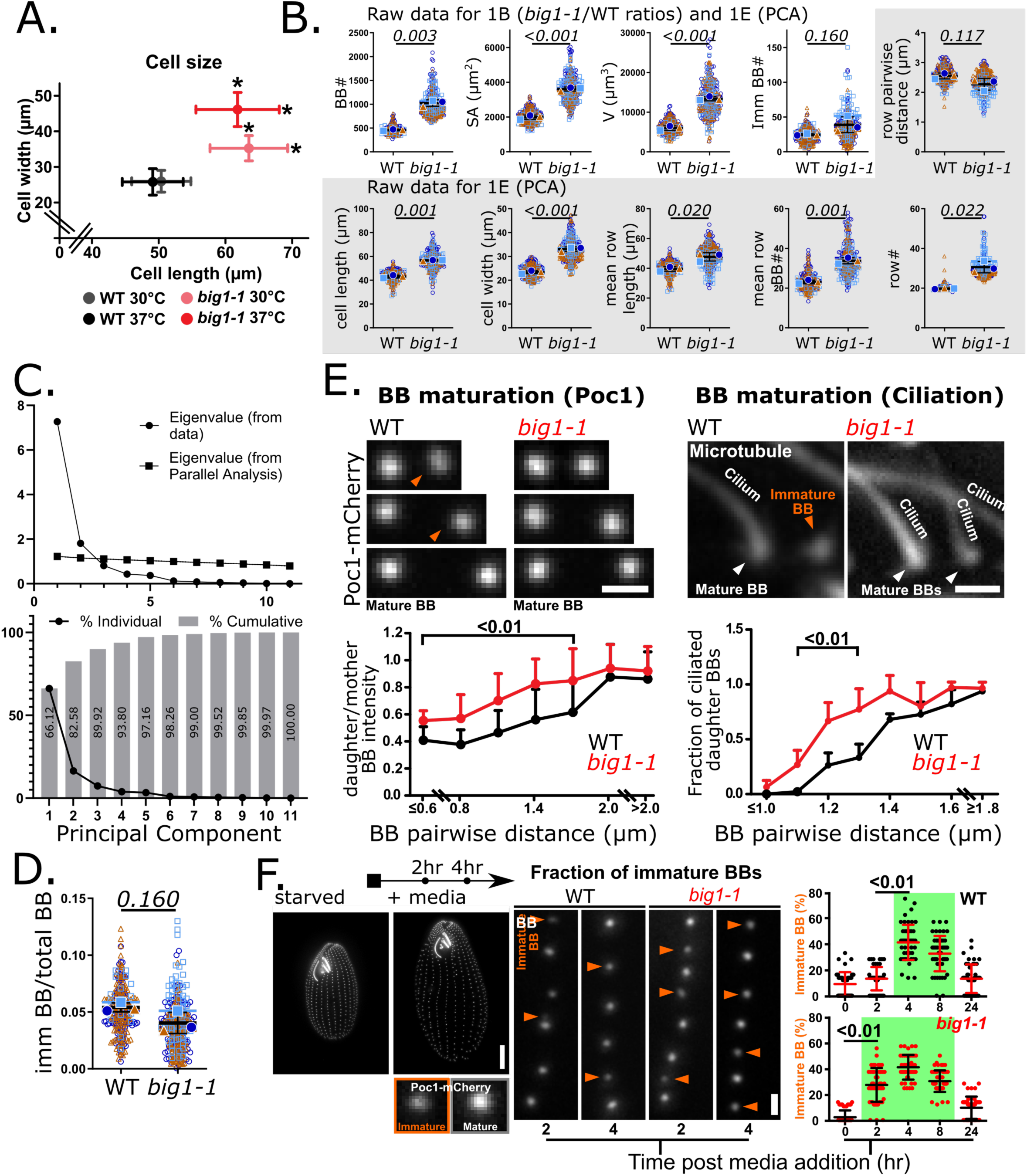
*big1-1* results in large cells with excess BBs that mature more rapidly than WT. A) Measurements of cell length and width at 30°C and 37°C for WT and *big1-*1 cells. Asterisks indicate p values <0.01. Error bars show standard deviations. B) Individual cell results supporting the averages of three biological replicates shown in Figure 1. Each symbol shows results from the automated image analysis for a single cell. The results reveal excess BBs, increases to surface area (SA), volume (V), length, width, cortical row length, BBs/row, cortical row number and the absolute number of immature BBs. Distance between cortical rows is reduced slightly. Error bars show standard deviations of the means of three biological replicates. C) PCA reveals that most variability is captured in PC1 and PC2. The percentage of variability captured by each PC is shown in the bottom graph. D) The ratio of imm BBs:total BB (data from A top left and fourth from the left) shows a slight reduction in imm BBs as a proportion of the total BBs in *big1-1*. Error bars show standard deviations of three biological replicates. E) BB daughter BB maturation occurs closer to mother BBs in *big1-1*. Left panels show Poc1-mCh intensity. Right panels show the ability to nucleate a cilium as a marker for daughter BB maturation. Graphs below show a significant decrease in distance from the mother in daughter BB maturation. Scale bar = 1 µm. Error bars show standard deviations. F) Cells arrested with mature BBs by starvation followed by release into media show *big1-1* cells build more BBs than WT cells. Dim Poc1 fluorescence intensities (red arrowheads) mark nascent BBs while bright Poc1 fluorescence intensities mark mature BBs. Scale bar = 10 µm for whole cell and 1 µm for insets. Graphs of the time course of the percentage of nascent BBs per 10 µm in the medial region of the cells. Error bars show standard deviations.

**Supplemental Figure 2.**
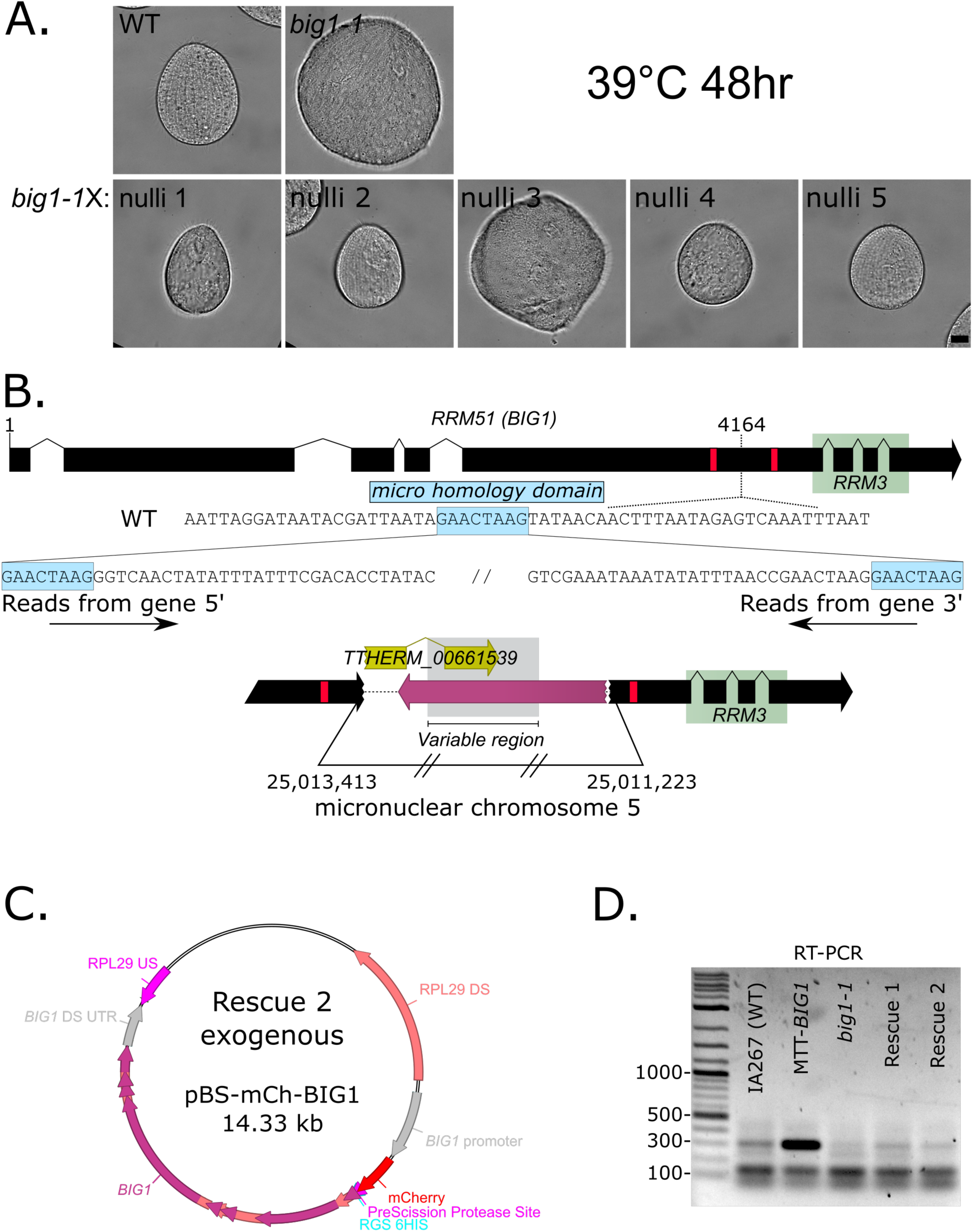
Insertion mutations in *RRM51* cause the *big1-1* phenotype. A) Top panels: DIC images of WT and *big1-1* cells grown at 39°C for 48 hours. Bottom panels show similarly grown cells after *big1-1* cells were crossed to nullisomic cell lines, indicating that the gene responsible for the *big1-1* phenotype is on MIC chromosome 3. Scale bar = 10 µm. B) A schematic of the insertion mutations in *RRM51/BIG1*. A microhomology domain appears to have been the site of insertional recombination as it appears on both ends of the inserted sequence. Sequencing of two of the prominent insertions reveals they include variable sequences from a single MIC chromosome 5 locus. C) A map of the exogenous Rescue2 construct showing the site of the mCh-6HIS tag. D) RT-PCR from RNA isolated from WT, *BIG1* overexpressing, *big1-1*, *big1-1*/Rescue1 and *big1-1*/Rescue2 cells confirming expression of the WT *BIG1* in rescued strains.

**Supplemental Figure 3.**
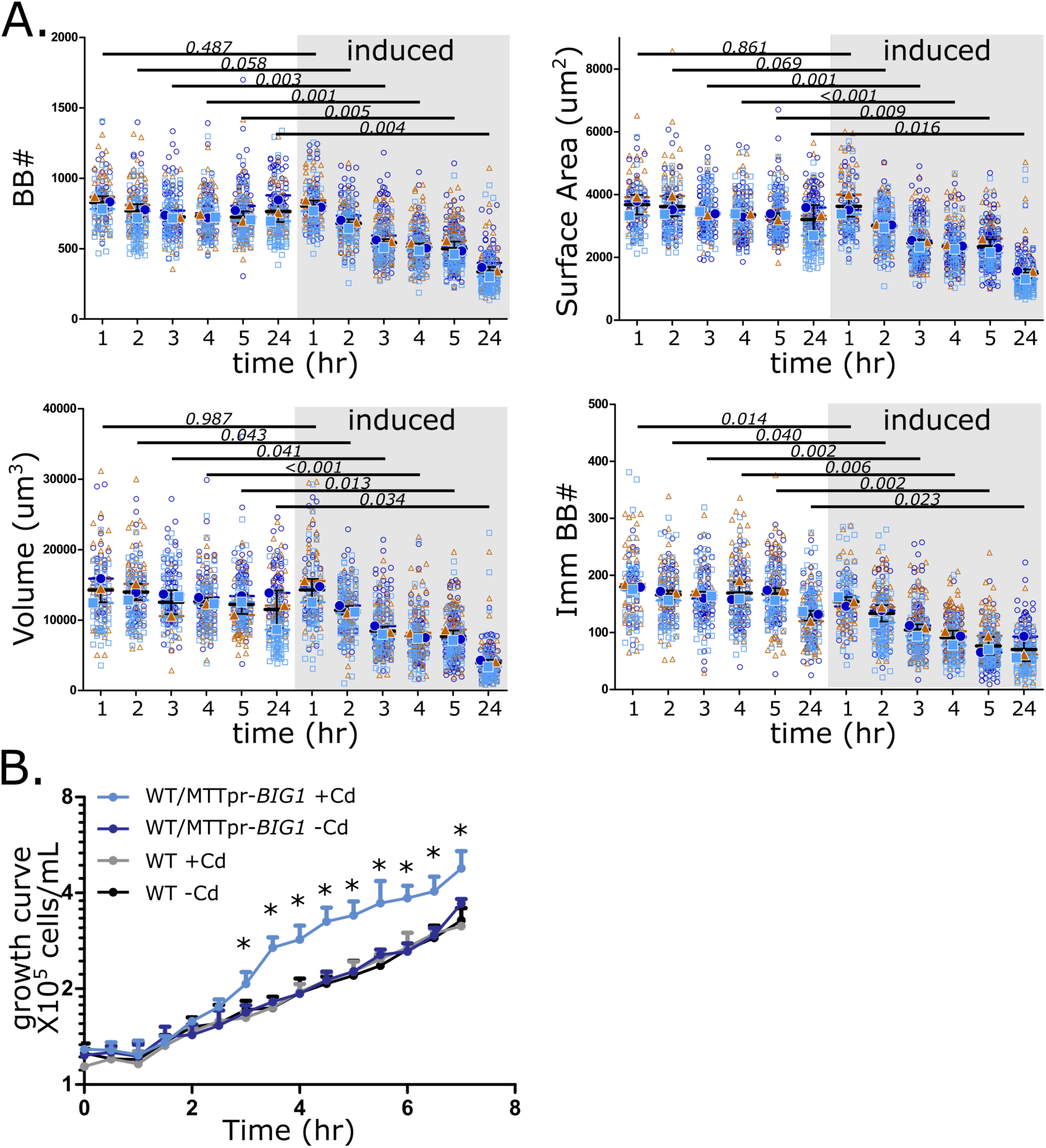
*BIG1* overexpression leads to small cells with fewer BBs. A) Quantifications from the automated image analysis over a time course of *BIG1* overexpression shows progressive loss of BBs, surface area, volume and immature BBs. B) Growth curves of WT and inducible *BIG1* expression cells (MTT1 promoter) in the presence or absence of the inducer (Cadmium). Counts were made every 30 minutes for seven hours. Asterisks indicate a p value <0.05. Error bars show the standard deviation of three biological replicates.

**Supplemental Figure 4.**
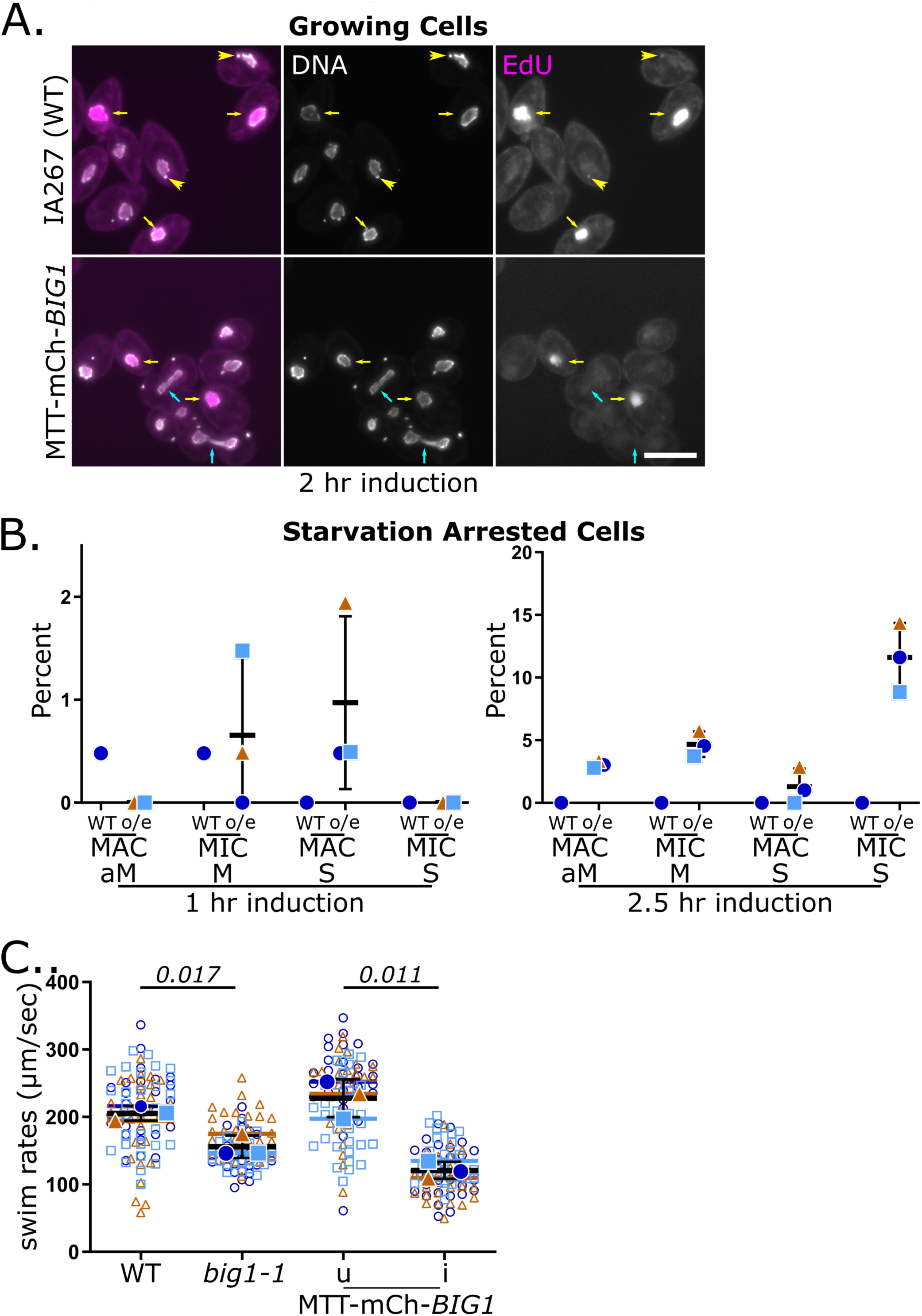
*BIG1-1* overexpression induces nuclear progression through the cell cycle and cell size optimizes swim rates. A) Two hours of *BIG1* overexpression in growing cells changes nuclear cell cycle stages. EdU staining shows cells in S phase. MAC S is marked by yellow arrows and MIC S by yellow arrowheads. Hoechst staining shows cells in MAC amitosis (cyan arrows), which happens just prior to the end of cell division. Scale bar = 50 µm. B) Nuclear stage quantification of starvation arrested cells after 1 or 2.5 hours of cadmium induced induction. One WT strain and three MTTpr-*BIG1* strains were measured. C) Swim speeds at 30°C of WT, *big1-1,* and MTTpr-*BIG1* cells with and without 24 hours of *BIG1* induction. Error bars show standard deviations of three biological replicates.

**Supplemental Table 1. Statistical methods.** The number of cells analyzed for each replicate are noted in black text. Blue text indicates which statistical tests were used. The table is organized by figure.

**Supplemental Table 2. Oligonucleotides used in this study.** Oligonucleotides are listed in the 5’ to 3’ direction with a brief description of the purpose of each one.

## Notes

### Competing Interest Statement

The authors have declared no competing interest.

